# RNA Sequencing in Adult *Drosophila* Females Identifies Estrogen-Related Receptor-Dependent Transcriptional Changes in Metabolism, DNA Replication, and Translation

**DOI:** 10.64898/2026.05.21.726871

**Authors:** Sophie A. Fleck, Eleanor B. Goldstone, Lesley N. Weaver

**Affiliations:** Department of Biology, Indiana University, Bloomington, IN 47405, USA

**Keywords:** nuclear receptor, glycolysis, pentose phosphate pathway, ribosome, translation

## Abstract

Nuclear receptors, transcription factors essential for organism growth, development, and reproduction, are expressed in a variety of tissues, with some exhibiting differential expression between males and females. The Estrogen-related receptor (ERR) is a conserved metabolic nuclear receptor required for energy metabolism and lipid accumulation. While previous studies in *Drosophila* have identified potential ERR targets from mixed sex larval populations and adult males, it is unclear whether transcriptional targets and biological pathways downstream of ERR are altered in a sex-specific manner. Here, we took an RNA sequencing approach to identify candidate ERR targets specifically in adult females and compared differentially expressed genes to a published male-specific dataset. Whole body conditional knockout of *ERR* significantly downregulated transcription of enzymes associated with glycolysis and the pentose phosphate pathway. In contrast, components of the DNA replication machinery were selectively downregulated in adult females, whereas ribosome biogenesis transcription was increased. Our results have further defined the metabolic targets of ERR between males and females, as well as suggest that ERR regulates DNA replication and global translation in females.

**SUMMARY:** In this manuscript, we used RNA sequencing to identify differential expression of transcripts dependent on the nuclear receptor ERR in *Drosophila* adult females. We find that ERR is required for activating transcription of glycolytic and pentose phosphate pathway enzymes, as observed in larvae and adult males. Furthermore, compared to males, loss of *ERR* in females specifically decreased DNA replication enzyme components while upregulating ribosomal components. Our results suggest that nuclear receptors have common and sex-specific targets, which will be of interest for those in the nuclear receptor and sexual dimorphism fields.

## INTRODUCTION

Maintenance of tissue homeostasis and energy metabolism requires precise coordination between organs (Droujinine and Perrimon 2016; Tokizane and Imai 2025). Members of the nuclear receptor superfamily behave as physiological sensors to alter the transcriptional landscape of the cell to promote development, cellular differentiation, immune activation, and reproduction (Mouzat *et al*. 2013; Fan and Evans 2015; Festuccia *et al*. 2017; Jin *et al*. 2025). In addition, nuclear receptors are regulators of sexual dimorphism, which allows for differential gene expression to support sex specific physiology (Rando and Wahli 2011; Dean *et al*. 2021; Xiong *et al*. 2023). For example, the mammalian Steroidogenic Factor-1 (SF-1, AD4BP) is required in the testes to promote expression of Müllerian Inhibiting Substance (MIS) for proper reproductive tract development (Giuili *et al*. 1997). Similarly, in *Drosophila* adults, the steroid hormone ecdysone is primarily produced in ovarian follicle cells to stimulate feeding behavior via Ecdysone Receptor signaling in the central nervous system, which in turn supports oogenesis (Sieber and Spradling 2015). However, the manner by which nuclear receptors dynamically alter their transcriptional targets during development and between the sexes has not been completely elucidated.

Estrogen-related receptors (ERRs) are conserved regulators of lipid and energy metabolism and are expressed in energy consuming tissues (Tennessen *et al*. 2011; Huss *et al*. 2015; Misra *et al*. 2017; BEEBE *et al*. 2020; Geng *et al*. 2024; Fasteen *et al*. 2025). In mammals, ERRα and ERRγ promote the transcription of enzymes involved in lipid metabolism and mitochondria production (Luo *et al*. 2003; Schreiber *et al*. 2004; Audet-Walsh and Giguere 2015; Fan and Evans 2015; Fan *et al*. 2018; Fox *et al*. 2022); whereas, ERRβ is required for maintaining stem cell pluripotency (Festuccia *et al*. 2012; Festuccia *et al*. 2017). The sole *Drosophila* ERR homolog is a central regulator of carbohydrate metabolism during development (Tennessen *et al*. 2011), promotes glucose oxidation and lipogenesis in adult males (Beebe *et al*. 2020), and is required for fertility in both adult males and females (Misra *et al*. 2017; Zike *et al*. 2025). Considering ERRs are essential metabolic regulators that can also control stem cell behavior, determining the conserved targets between organisms and the sexes may provide insight to possible targets for therapeutic intervention.

In this study, we used RNA sequencing of *ERR* conditional knockout alleles in *Drosophila* adult females to refine the potential ERR targets that are conserved throughout development and between the sexes. Using two different *ERR* conditional alleles, we identified a core set of glycolytic and pentose phosphate pathway (PPP) enzymes that require ERR for their transcription. Furthermore, by comparing our data to a published adult male-specific dataset, we determined that regulation of glycolysis and the PPP is a conserved feature of ERR. Lastly, we determined that ERR is required in females to support DNA replication and refine global translation. Our results suggest that ERRs are a key regulator of metabolism across species and has sex-specific transcriptional targets in *Drosophila* females.

## MATERIALS AND METHODS

### Drosophila strains and culture

*Drosophila* strains were maintained on BDSC cornmeal food (15.9 g/L inactive yeast, 9.2 g/L soy flour, 67.1 g/L yellow cornmeal, 5.3 g/L agar, 70.6 g/L light corn syrup, 0.059 M propionic acid) at 22-25^°^C. All females used in experiments were maintained on BDSC cornmeal food supplemented with wet active yeast paste.

The previously described *ERR* allele were used, including *dERR*^*Cond*.*19-4*^ [RRID:BDSC_91656, (Beebe *et al*. 2020)], *dERR*^*cond*.*2*^ (Beebe *et al*. 2020), *ERR*^*1*^ [RRID:BDSC_83688, (Tennessen *et al*. 2011)], and *ERR*^*2*^ [RRID:BDSC_83689, (Tennessen *et al*. 2011)]. In the *ERR*^*cond*.*19-4*^ construct, the endogenous *ERR* locus is flanked by two FRTs and is in the background of the previously described *ERR*^*2*^ allele (Tennessen *et al*. 2011), which is a deletion of the entire *ERR* coding region plus portions of *atg18* and *CG7979*. Heat shock removes the majority of the *ERR* coding region, generating an *ERR* null animal. For the *ERR*^*cond*.*2*^ construct, two FRTs surround a fluorescently tagged *ERR* BAC transgene (*ERR::GFP*.*FSFTF*) that is in trans to *ERR*^*1*^ and *ERR*^*2*^ mutant alleles. The previously described *ERR*^*1*^ allele contains an 84 bp deletion that removes the exon 2 splice acceptor, as well as two additional point mutations in exons 2 and 3 (Tennessen *et al*. 2011). Upon heat shock, an *ERR*^*1/2*^ mutant animal is generated. Animals carrying these constructs develop normally prior to adulthood when *ERR* knockout is induced by heat shock as previously described (Beebe *et al*. 2020; Zike *et al*. 2025).

Flybase (flybase.org) was used as a reference tool throughout this study (Ozturk-Colak *et al*. 2024).

### RNA isolation

Ten adult females of each control or *ERR* conditional knockout genotype were flash frozen in liquid nitrogen and lysed in 500 µl lysis buffer from the RNAqueous-4PCR DNA-free RNA isolation for RT-PCR kit (Invitrogen) as previously described (Zike *et al*. 2025). RNA was extracted from three independent experiments using manufacturer’s instructions.

### RNA sequencing (RNA-seq)

cDNA library preparation, Illumina sequencing, and differential expression analysis was performed by Novogene Bioinformatics Technology Co., Ltd (Beijing, China). cDNA was prepared using the NEBNext Ultra RNA Library Prep Kit for Illumina (New England Biolabs) according to manufacturer’s instructions. Quality assessment for the cDNA library for each sample was assessed using an Aligent Bioanalyzer 2100, followed by sequencing on a NovaSeq6000 platform with PE150 read lengths.

Sequencing reads were aligned to the *D. melanogaster* reference genome using the TopHat read alignment tool (Trapnell *et al*. 2009). Reference sequences were downloaded from the Ensembl project website (useast.ensembl.org). TopHat alignments generated read counts for each gene using HTSeq, which were subsequently used to generate differential expression results using the DESeq2 R package (Anders *et al*. 2015). Differentially expressed genes with a corrected *P* value less than 0.5 were considered significant. Gene ontology (GO), KEGG pathway, and REACTOME pathway enrichment of differentially expressed genes were analyzed using PAthway, Network and Gene-set Enrichment Analysis (PANGEA) (Hu *et al*. 2023).

## RESULTS AND DISCUSSION

### RNA-seq of ERR conditional knockout females to identify differentially expressed genes

ERR-dependent transcripts have been identified in mixed-sex larvae (Tennessen *et al*. 2011) and adult males (Beebe *et al*. 2020); however, an adult female-specific dataset is currently unavailable. To identify downstream transcriptional targets of ERR specifically in adult females, we performed RNA-seq analysis from control and whole body *ERR* conditional knockout females, which allows animals to develop normally prior to adulthood when *ERR* deletion is induced. We induced sequential heat shock treatments at 37^°^C (separated by 24 hours) to zero-to-two day old *ERR*^*cond*.*19-4*^ or *ERR*^*cond*.*2*^ females and compared the differentially expressed genes to their respective no heat shock sibling control 7 days after heat shock (AHS). We and others previously showed that this manipulation and time course is sufficient to downregulate known ERR targets (Beebe *et al*. 2020; Zike *et al*. 2025).

RNA-seq produced an average of 45,427,224 reads across the twelve sequencing libraries (each respective control and *ERR* conditional knockout condition in triplicate), mapping an average 94% of the reads to the *Drosophila melanogaster* genome (Adams *et al*. 2000) (**Table S1**). Over 14,000 transcripts were mapped to protein coding genes, whereas about 1,000 transcripts were identified as either pseudogenes, ribosomal RNA (rRNA), small nucleolar RNA (snoRNA), small nuclear RNA (snRNA), or transfer RNA (tRNA) (**Tables S2 and S3**). Although we enriched for mRNAs using oligo(dT) beads for poly(A) selection, this method does not eliminate non-coding RNAs that contain poly(A) tails (Cai *et al*. 2004; Wilhelm *et al*. 2008; Cabili *et al*. 2011).

*ERR* knockout using the *ERR*^*cond*.*2*^ allele identified 557 significantly differentially expressed transcripts (padj ≤ 0.05, **Figure 1A**), with most transcripts representing protein coding genes. Of those differentially expressed transcripts, 285 were downregulated protein coding genes (indicating positive regulation by ERR), and ~260 were significantly upregulated (indicating genes negatively regulated by ERR activity) (**Figure 1C**). In the *ERR*^*cond*.*19-4*^ condition, ~400 genes were differentially regulated (**Figure 1B**), with 111 protein coding genes significantly downregulated and 288 transcripts upregulated (**Figure 1D**).

**Figure 1.**
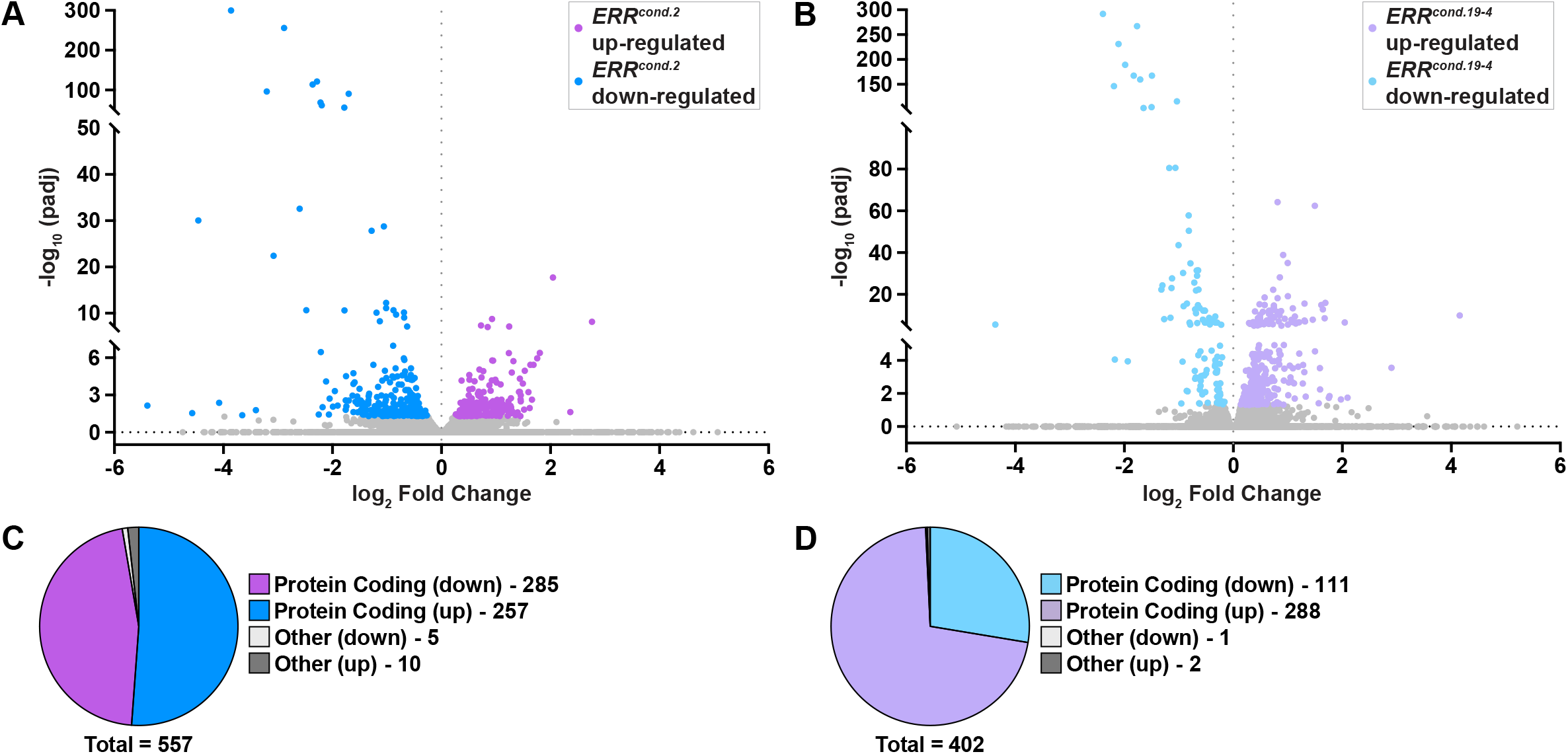
Conditional knockout of *ERR* in adult females differentially regulates over 400 genes. **(A, B)** Volcano plots of differentially expressed genes from *ERR*^*cond*.*2*^ **(A)** or *ERR*^*cond*.*19-4*^ **(B)** females graphing the statistical significance [-log_10_(padj)] to the magnitude of differential expression (log_2_ Fold Change). **(C, D)** Pie charts comparing the number of upregulated and downregulated genes in *ERR*^*cond*.*2*^ **(C)** or *ERR*^*cond*.*19-4*^ **(D)** females.

The difference in differentially expressed transcripts is most likely due to differences in allelic strength. We previously reported that the *ERR*^*cond*.*2*^ allele significantly decreased known components of the pentose phosphate pathway (PPP) compared to the weaker *ERR*^*cond*.*19-4*^ allele (Zike *et al*. 2025), suggesting the *ERR*^*cond*.*2*^ allele may be a stronger manipulation. Collectively, our results suggest that our transcriptomic approach identifies the majority of genes expressed in adult *Drosophila melanogaster* females and identifies significantly differentially expressed genes in the absence of *ERR*.

### ERR knockout in adult females downregulates the expression of 117 transcripts

To identify the candidates most likely to be direct targets of ERR in adult females, we first compared the differentially expressed genes that were downregulated in the *ERR*^*cond*.*2*^ and *ERR*^*cond*.*19-4*^ alleles. Between the two conditions, 49 common transcripts were significantly decreased, where 241 downregulated transcripts (combining “protein coding” and “other”) were enriched in the *ERR*^*cond*.*2*^ females compared to 63 unique transcripts in *ERR*^*cond*.*19-4*^ females (**Figure 2A**). To determine whether these genes clustered in similar biological pathways or enzymatic function, we performed enrichment analysis of our gene set using PAthway, Network, and Gene-set Enrichment Analysis (PANGEA) (Hu *et al*. 2023). Genes involved in metabolic pathways, carbohydrate metabolism, the PPP, and glycolysis were enriched in the most statistically significant categories (**Table S4**). In both conditions, transcript levels for components of glycolysis and the PPP were less abundant in the absence of *ERR* (**Figure 2B-H**). For example, transcripts for the glycolytic enzymes *Gapdh1, Gaphd2*, and *Tpi* were significantly decreased in both *ERR*^*cond*.*2*^ and *ERR*^*cond*.*19-4*^ females (**Figure 2C-E**). Furthermore, known PPP component transcripts (*Zw, Rpi*, and *Pgd*) were also significantly decreased when *ERR* was conditionally knocked out in adult females (**Figure 2F-H**). Consistent with previous reports (Tennessen *et al*. 2011; Beebe *et al*. 2020; Fasteen *et al*. 2025), our results confirm that ERR is a conserved central regulator of metabolism between organisms, at different developmental stages, and between the sexes.

**Figure 2.**
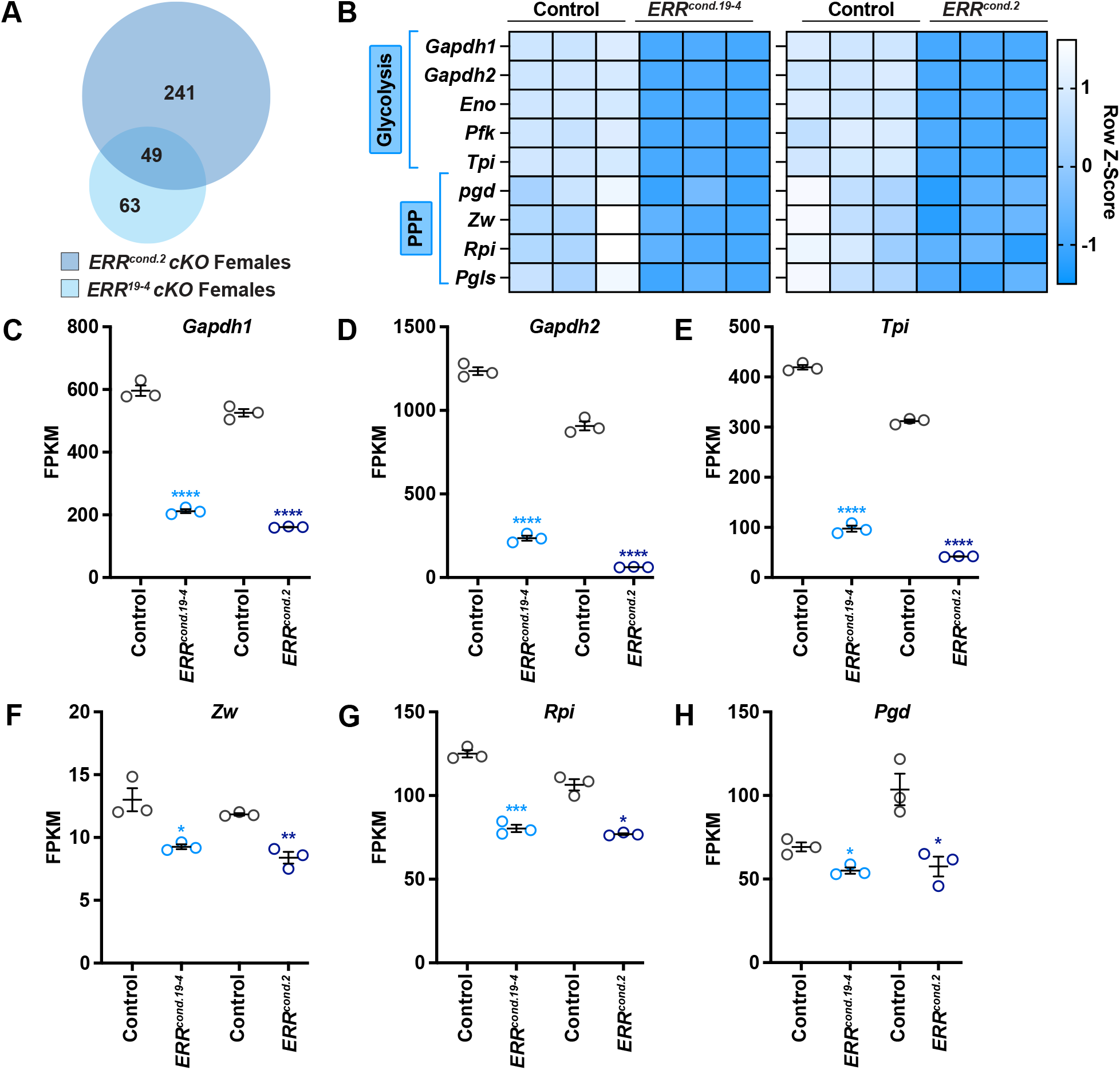
Loss of *ERR* in adult females downregulates glycolytic and Pentose Phosphate Pathway enzymes. **(A)** Venn diagram indicating the common and distinct downregulated transcripts in *ERR*^*cond*.*2*^ and *ERR*^*cond*.*19-4*^ females. **(B)** Heatmap showing the relative glycolytic and pentose phosphate pathway transcript levels in control, *ERR*^*cond*.*2*^, and *ERR*^*cond*.*19-4*^ females. **(C-H)** Quantification of normalized transcript expression (FPKM) for glycolytic **(C-E)** and pentose phosphate pathway **(F-H)** enzymes in control, *ERR*^*cond*.*2*^, and *ERR*^*cond*.*19-4*^ adult females. Data shown as mean ± SEM. \*\*\*\**P* < 0.0001, \*\**P* < 0.01, \**P* < 0.05.

### Loss of ERR in adult females upregulates fatty acid metabolism genes

Loss of *ERR* also resulted in transcriptional upregulation, suggesting possible indirect mechanisms of transcript repression under normal conditions. Inducing *ERR* conditional knockout with both alleles significantly upregulated 28 common transcripts (**Figure 3A**). We next used PANGEA to determine whether these upregulated genes clustered according to biological function. PANGEA analysis (**Table S5**) showed that enzymes required for fatty acid metabolism were enriched in the common upregulated genes. For example, in both *ERR* conditional knockout alleles, *Acyl-CoA binding protein 3* (*Acbp3*) and *CG16985* transcripts were significantly increased (**Figure 3B and C**). Members of the ACBP family prevent acyl-CoA hydrolysis to regulate lipid metabolism (Majerowicz *et al*. 2016); whereas *CG16985* encodes a predicted fatty acyl-CoA hydrolase, which hydrolyzes fatty acyl-CoAs into free fatty acids and CoA to control peroxisomal and mitochondrial oxidation (Caswell *et al*. 2022). Although not significant for *ERR*^*cond*.*19-4*^ the fatty acid synthase, *FASN1*, was also upregulated (**Figure 3D**). These results suggest that in the absence of *ERR*, adult females upregulate fatty acid metabolism enzymes to modulate energy production.

**Figure 3.**
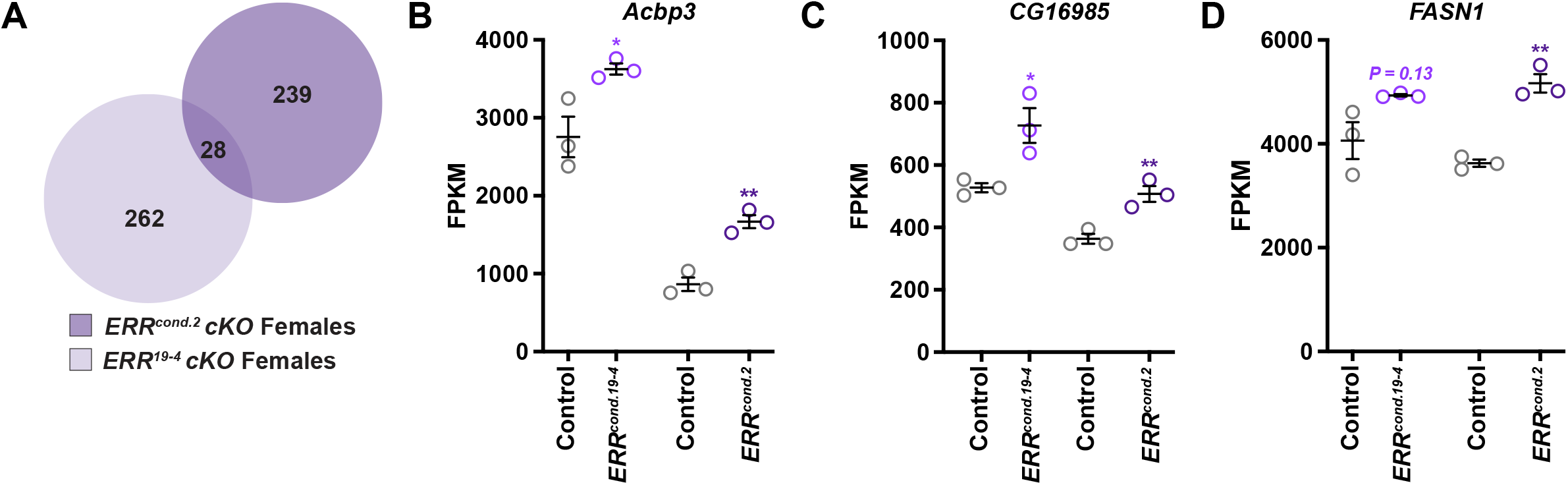
Enzymes involved in fatty acid synthesis are upregulated in the absence of *ERR*. **(A)** Venn diagram showing the common overlap of transcripts significantly upregulated in *ERR*^*cond*.*2*^ and *ERR*^*cond*.*19-4*^ adult females. **(B-D)** Quantification of normalized transcript expression (FPKM) for fatty acid metabolism genes. Data shown as mean ± SEM. \*\**P* < 0.01, \**P* < 0.05.

ERRs are regulators of mitochondrial energy metabolism and have been shown to positively regulate the transcription of oxidative phosphorylation and fatty acid oxidation enzymes (Cartoni *et al*. 2005; Rangwala *et al*. 2010; Fan and Evans 2015; Fan *et al*. 2018; Fox *et al*. 2022; Ma *et al*. 2025). Furthermore, mammalian ERRs are upregulated in energetically demanding tissues under stress. For example, short-term exercise induces transcription of *ERRα* and *ERRγ* in skeletal muscle (Wende *et al*. 2005) and *ERRα* null myotubules have reduced fatty acid oxidation capacity (Dufour *et al*. 2007). In *Drosophila* larval adipocytes, *ERR* mutants have decreased lipid accumulation due to decreased expression of fatty acid β-oxidation genes (Fasteen *et al*. 2025). Collectively, these results suggest that ERR is required to maintain energy metabolism via regulation of fatty acid metabolism by promoting synthesis and/or catabolism in a tissue-specific manner.

### Whole body loss of ERR in adult females decreases DNA replication machinery transcription

Metabolism across species is sexually dimorphic (Shingleton and Vea 2023; Mauvais-Jarvis 2024), and differences in nuclear receptor signaling between the sexes has been implicated in physiological changes (Dean *et al*. 2021). We therefore compared our adult female dataset to a published adult male specific dataset in which *ERR* was conditionally knocked using the *ERR*^*cond*.*2*^ allele (Beebe *et al*. 2020). Loss of *ERR* in adult males and females decreased the transcription of 97 common genes; whereas 193 genes were uniquely downregulated in females compared to 650 unique transcripts in males (**Figure 4A, Table S6**). We speculate that the significant difference in the number of differentially expression genes between the sexes is due to *ERR* knockout during pupariation for the male dataset (Beebe *et al*. 2020), compared to adult-specific knockout for females. Analysis of the common downregulated genes using PANGEA showed that these transcripts clustered in genes involved in metabolic pathways including carbon metabolism and the PPP (**Figure 4B, Table S7**). We next used PANGEA to determine the biological pathways that were specifically downregulated in adult females. Decreased genes specifically in *ERR* conditional knockout females were involved in DNA replication (**Figure 4C, Table S8**). *PolD1* (a subunit of DNA polymerase δ), *Mcm3* (a component of the DNA helicase complex), and *Orc2* (a component of the origin of recognition complex) transcripts were significantly downregulated in *ERR*^*cond*.*2*^ adult females (**Figure 4D**).

**Figure 4.**
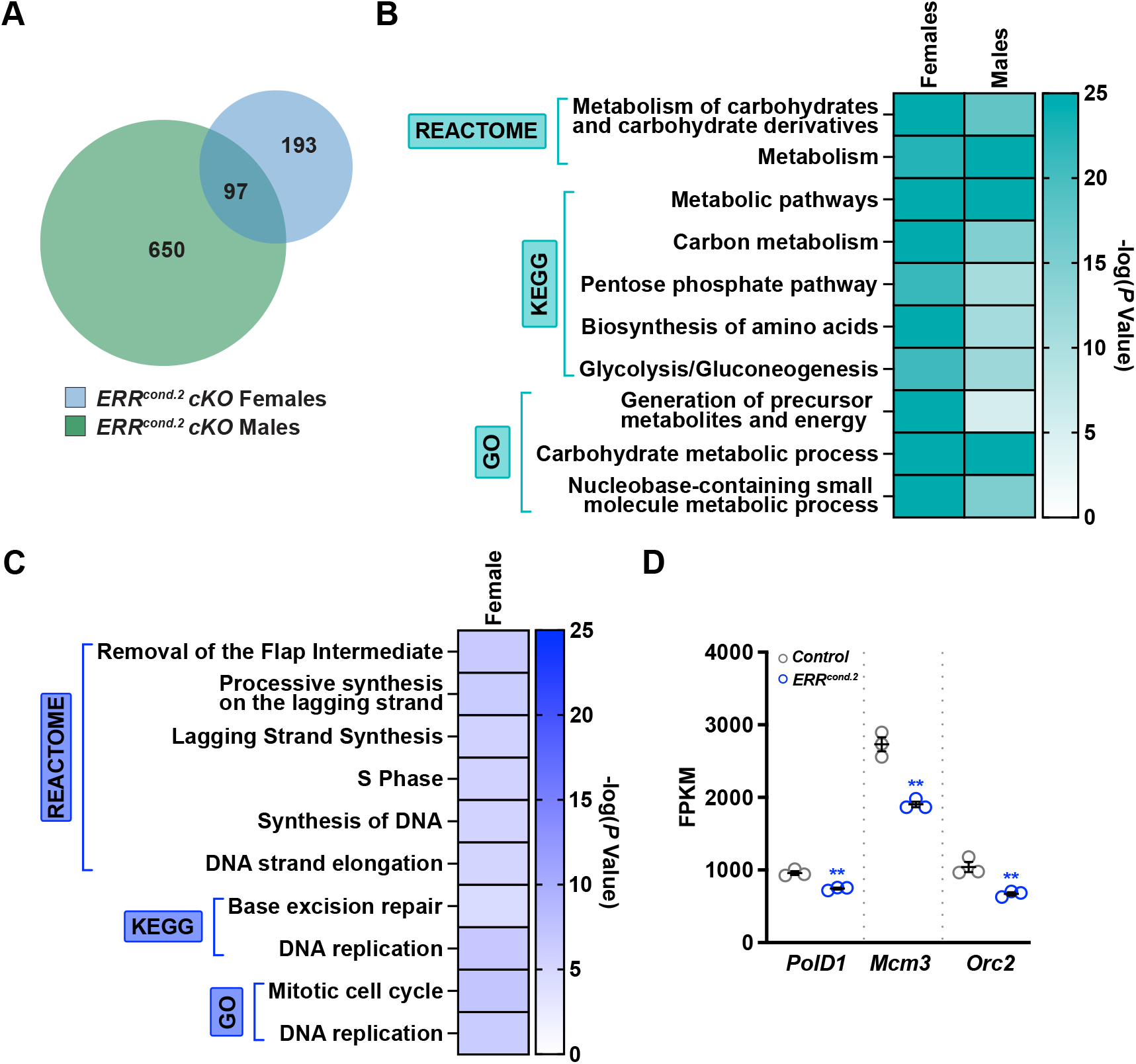
Downregulation of DNA replication machinery components are enriched in *ERR* conditional knockout females. **(A)** Venn diagram indicating the overlap of significantly decreased transcripts in adult *ERR*^*cond*.*2*^ males and females. **(B)** PANGEA enrichment analysis of differentially expressed genes in males and females in the absence of *ERR*. Heatmap indicates the enriched gene sets based on the rank-ordered adjusted *P*-value. **(C)** Heatmap indicating the top 10 enriched PANGEA gene sets specifically downregulated in adult *ERR*^*cond*.*2*^ females based on the adjusted *P*-value. **(D)** Quantification of normalized transcript expression (FPKM) for DNA replication machinery. Data shown as mean ± SEM. \*\**P* < 0.01.

One hypothesis for the decreased expression of DNA replication enzymes in females lacking *ERR* is to halt oogenesis, an energetically demanding process (Drummond-Barbosa and Spradling 2001; Laws and Drummond-Barbosa 2017). At mid-oogenesis follicle cells in the developing egg chamber switch from mitotic to endocycling cells in which the follicle cells become polyploid by repeatedly duplicating the genome without dividing (Calvi *et al*. 1998). Because the ovary responds to a variety of physiological cues (Drummond-Barbosa 2019) and is regulated by nuclear receptor-mediated inter-organ communication (Weaver and Drummond-Barbosa 2019; Weaver and Drummond-Barbosa 2021), whole body loss of *ERR* in females may indirectly downregulate DNA replication in tissues with polyploid cells, such as the ovary.

### Ribosome biogenesis is selectively increased in ERR conditional knockout females

Comparison of the upregulated genes between *ERR* conditional knockout males and females showed that only 16 transcripts were common between the sexes; where 240 transcripts were enriched in males, and 251 transcripts were selectively upregulated in females (**Figure 5A, Table S9**). PANGEA analysis of the 16 commonly upregulated transcripts suggested enzymes involved in oxidative phosphorylation and aerobic respiration are altered in the absence of *ERR*, however, these processes were mostly enriched in the male dataset (**Figure 5B**). Analysis of the female-specific transcripts showed an upregulation in genes that promote translation (**Figure 5C, Table S10**). Structural components of the ribosome were significantly upregulated in *ERR*^*cond*.*2*^ females (**Figure 3F-H**), suggesting defects in ERR signaling may activate global translation in adult females. Interestingly, the mammalian ortholog ERRα has been shown to selectively activate transcript expression of *Rplp1* (ribosomal protein lateral stalk subunit P1) and genes involved in autophagy in response to starvation (Tripathi *et al*. 2025), suggesting ERRα is required for stress adaptation. In contrast, our results suggest that signaling downstream of ERR restricts overall translation.

**Figure 5.**
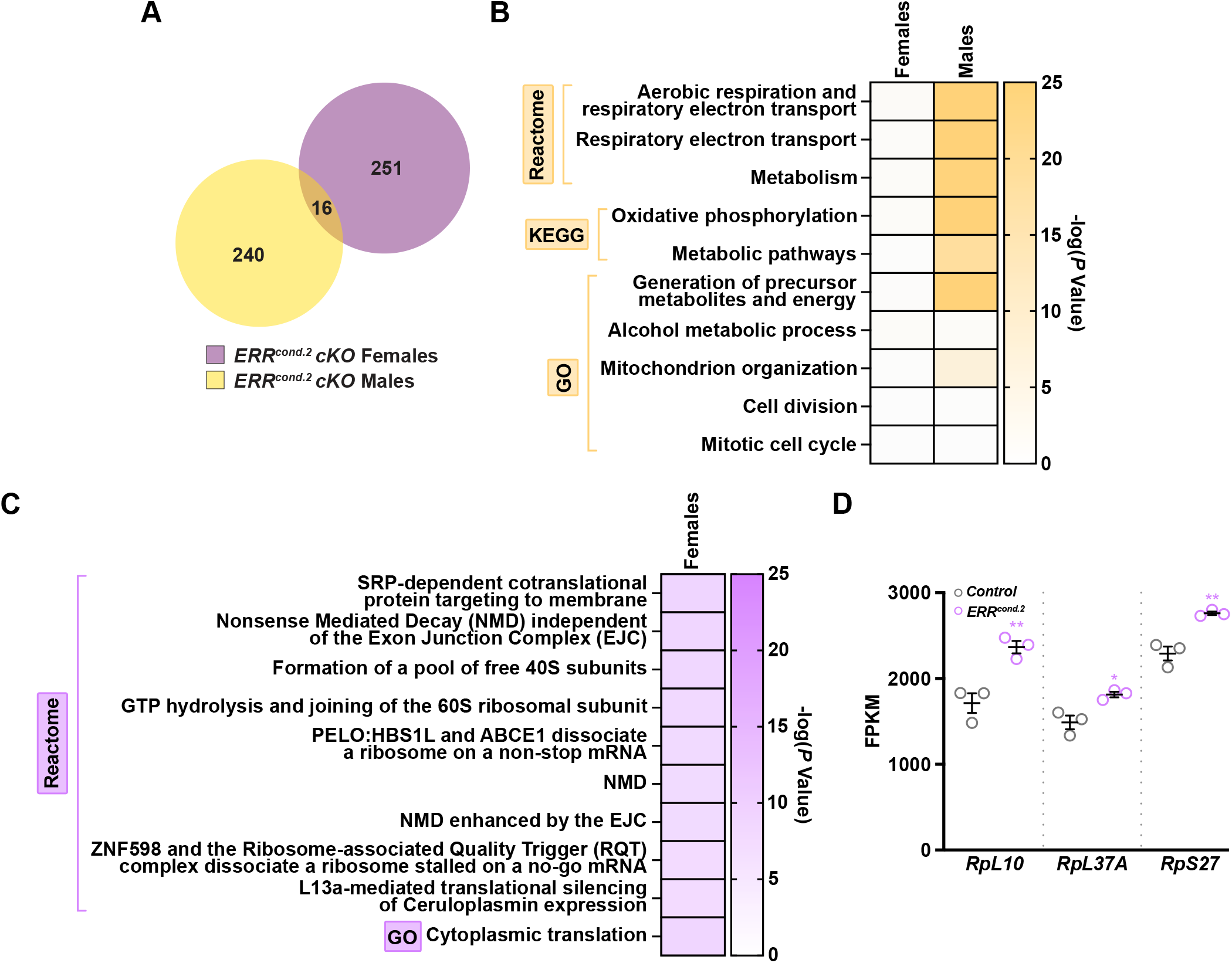
Ribosome biogenesis components are specifically upregulated in *ERR* conditional knockout females. **(A)** Venn diagram showing the overlap of significantly upregulated transcripts in *ERR*^*cond*.*2*^ adult males and females. **(B)** PANGEA enrichment analysis of differentially expressed genes in males and females lacking *ERR*. Heatmap indicates enriched gene sets based on the adjusted *P*-value. **(C)** Heatmap indicating the top 10 enriched PANGEA gene sets specifically upregulated in *ERR*^*cond*.*2*^ adult females based on the adjusted *P*-value. **(D)** Quantification of normalized transcript expression (FPKM) for ribosome components. Data shown as mean ± SEM. \*\**P* < 0.01, \**P* < 0.05.

Collectively, our RNA sequencing analysis suggests that ERRs are conserved metabolic regulators in eukaryotes by promoting the transcription of glycolytic and PPP enzymes. Our analysis also suggests that in *Drosophila* there are sex-specific changes in the transcriptome in the absence of *ERR* most likely due to the energetic demand of the ovary. One limitation of our study is that our analysis was performed on whole-body samples in which *ERR* was knocked out in every tissue. Future studies should address whether direct targets of ERR are altered in different tissues and between the sexes.

## DATA AVAILABILITY

*Drosophila* strains are available from the Bloomington *Drosophila* Stock Center (BDSC). The data and analyses in this paper are described in the main. The raw data and processed data files for the adult female samples are available through the NCBI GEO accession number GSE331492 and are also provided as supplemental tables.

## ACKNOWLEDGMENTS

Stocks obtained from the BDSC (NIH P40OD018537) were used in this study. We are thankful to Flybase (flybase.org, NIH 5U24HG013300), an invaluable *Drosophila* research resource. We are grateful to members of the Weaver Lab and Alissa Armstrong for critical reading of the manuscript. This work was supported by National Institutes of Health R00 GM127605 and R35 GM150517 grants to L.N.W.

## FUNDING

This work was supported by National Institutes of Health R00 GM127605 and R35 GM150517 grants to L.N.W.

## CONFLICT OF INTEREST

The authors declare no conflicts of interest.

## SUPPLEMENTARY TABLES

**Table S1. Total number of mapped reads for each *ERR* conditional knockout library**.

**Table S2. List of all transcripts and their relative abundance in control and *ERR***^***cond*.*2***^ **females**.

**Table S3. List of all transcripts and their relative abundance in control and *ERR***^***cond*.*19-4***^ **females**.

**Table S4. PANGEA analysis of common downregulated genes in *ERR* conditional knockout females**.

**Table S5. PANGEA analysis of common upregulated genes in *ERR* conditional knockout females**.

**Table S6. Common and sex-specific genes downregulated in *ERR***^***cond*.*2***^ **conditional knockouts**.

**Table S7. PANGEA analysis of common downregulated genes in *ERR***^***cond*.*2***^ **conditional knockout females and males**.

**Table S8. PANGEA analysis of downregulated genes specifically in *ERR***^***cond*.*2***^ **females**.

**Table S9. Common and sex-specific genes upregulated in *ERR***^***cond*.*2***^ **conditional knockouts**.

**Table S10. PANGEA analysis of upregulated genes specifically in *ERR***^***cond*.*2***^ **females**.

## Notes

### Competing Interest Statement

The authors have declared no competing interest.

